# LRRC33 is a novel binding and regulating protein of TGF-β1 function in human acute myeloid leukemia cells

**DOI:** 10.1101/560763

**Authors:** Wenjiang Ma, Yan Qin, Bjoern Chapuy, Chafen Lu

## Abstract

Transforming growth factor – β1 (TGF-β1) is a versatile cytokine. It has context-dependent pro- and anti-cell proliferation functions. Activation of latent TGF-β1 requires release of the growth factor from pro-complexes and is regulated through TGF-β binding proteins. Two types of TGF-β binding partners, latent TGF-β-binding proteins (LTBPs) and leucine-rich-repeat-containing protein 32 (LRRC32), have been identified and their expression are cell specific. TGF-β1 also plays important roles in acute myeloid leukemia (AML) cells. However, the expression of LTBPs and LRRC32 are lacking in myeloid lineage cells and the binding protein of TGF-β1 in these cells are unknown. Here we show that a novel leucine-rich-repeat-containing protein family member, LRRC33, with high mRNA level in AML cells, to be the binding and regulating protein of TGF-β1 in AML cells. Using two representative cell lines MV4-11 and AML193, we demonstrate that the protein expression of LRRC33 and TGF-β1 are correlated. LRRC33 co-localizes and forms complex with latent TGF-β1 protein on the cell surface and intracellularly in these cells. Similar as in other cell types, the activation of TGF-β1 in MV4-11 and AML193 cells are also integrin dependent. We anticipate our study to be a starting point of more comprehensive research on LRRC33 as novel TGF-β regulating protein and potential non-genomic based drug target for AML and other myeloid malignancy.

## Introduction

Transforming growth factor – β1 (TGF-β1) is the primary member of the large transforming growth factor-β (TGF-β) family which have crucial roles in multiple processes including cell proliferation, development, wound healing and immune responses (*1, 2*). Abnormality of TGF-β function has been implicated in multiple human diseases, including fibrosis, autoimmune diseases and cancer (*3*). TGF-β1 is synthesized and secreted in a latent, inactive complex, which contains dimerized non-covalently associated TGF-β1growth factor domain and a large prodomain, the latency associated peptide (LAP) (*4*). Throughout this paper we use pro-TGF-β1 to indicate the furin-cleaved latent TGFβ protein. The pro-TGF-β1 latent protein does not have biological activity, thus the release of active TGF-β1 is a critical step for regulating TGF-β1 function in cell signaling.

The activation of the latent TGF-β1 is orchestrated by its binding proteins (*5*). There are several known binding partners of pro-TGF-β1. The latent transforming growth factor β binding proteins (LTBPs) consist of 4 isoforms (LTBP-1, -2, -3, and -4), that forms latent complexes with pro-TGF-β1 by covalently binding to LAP via disulfide bonds (*6-8*). LTBP is important in the assembly, storage, and secretion of TGF-β1 in that it targets pro-TGF-β1 to the extracellular matrix and leads to the release of soluble active TGF-β1 upon integrin dependent signaling pathways (*5*). Unlike LTBPs that associate with pro-TGF-β1 in extracellular matrix, another protein, glycoprotein-A repetitions predominant protein (GARP), also known as leucine rich repeat containing protein 32 (LRRC32), is a cell membrane associated protein that binds to LAP and directs pro-TGF-β1 to the cell surface of FOXP3+ regulatory T cells. The GARP-pro-TGF-β1 complex are stored on the cell surface and the integrin-dependent signaling pathway is also required for the release of active TGF-β1 (*9-11*).

TGF-β1 protein is pleiotropic in regulating all stages of hematopoiesis and it has both proliferative and anti-proliferative effects on different cells specific to cell types and cell differentiation stages (*12, 13*). Thus, TGF-β1 and its binding proteins have long been potential targets of therapies for different blood cancers. It has been reported that in multiple human acute myeloid leukemia (AML) cell lines, including OCI-AML1, AML193, and THP-1 cells, there are TGF-β1 expression, and the proliferation and differentiation of these cells are affected by TGF-β1 through autocrine and paracrine pathways (*14, 15*). However, the regulation of TGF-β1 activation in myeloid leukemia cells is not clearly understood. Previous studies show that LTBPs are expressed primarily in cell types of mesenchymal origin (*16*) and LRRC32 is reported to mainly express on endothelium cells, platelets, and Foxp3+ regulatory T cells but not on myeloid cells (*17*). We recently reported that LRRC33, a homologous protein of the pro-TGF-β1 binding protein GARP (LRRC32), is covalently linked to the prodomain of TGF-β1, and highly expressed microglia cells in the central nervous system (CNS) where LRRC33 associates with pro-TGF-b1 and regulates TGF-β1 function (*18*). Thus, LRRC33 is the potential binding partner of pro-TGF-β1 in other myeloid cells, including human AML cells.

In this study, we showed that LRRC33 and pro-TGF-β1 co-localize and form a protein complex through disulfide bonds on the cell surface of two human acute myeloid leukemia cell lines: MV4-11 and AML193. We show that the activation of TGF-β1 in MV4-11 and AML193 cells is α_V_ integrin–dependent and correlated with the expression level of LRRC33. Our results suggest that LRRC33 plays an important role in the regulation of TGF-β1 activation in acute myeloid leukemia cells.

## Methods and Materials

### The mRNA expression of LRRC33 and TGF-β1

The mRNA expression data is obtained by Affymetrix Arrays from Cancer Cell Line Encyclopedia of Broad Institute (Boston, MA). The raw data are analyzed and plotted using Prism 7 Graphpad software.

### Cell culture and Phorbol 12-myristate 13-acetate (PMA) stimulation of cells

MV4-11 and AML193 cells were obtained from ATCC and were cultured in Iscove’s Modified Dulbecco’s Medium (IMDM) (ThermoFisher Scientific, Waltham, MA) supplemented with 10% fetal bovine serum (FBS), 5 mM L-glutamine, 1% nonessential amino acids, and 5mM penicillin/streptomycin. Culture medium for AML193 cells was supplemented with 0.005 mg/ml insulin and 5 ng/ml GM-CSF (Sigma-Aldrich, St. Louis, MO). Cells were cultured at 37°C in a humidified 5% CO2 atmosphere. Phorbol 12-myristate 13-acetate (PMA) (Sigma-Aldrich, St. Louis, MO) was dissolved in DMSO at the concentration of 1.6 mM/ml, then diluted to 80 nM in cell culture media and incubated with MV4-11 or AML193 cells for 24 hours for cell stimulation.

### Antibodies and other reagents

The following antibodies were used in this study: proTGFβ1 antibody TW7-28G11 for Immunoprecipitation (IP) and TW4-2F8 for flow cytometry (*18*), biotinylated anti-LAP1 antibody for Western blot (WB) (BAF246; R&D Systems), polyclonal goat anti-LAP1 antibody (R&D Systems), APC-conjugated anti-proTGFβ1 (R&D Systems), mouse IgG (ThermoFisher Scientific), goat IgG (ThermoFisher Scientific) for flow cytometry and confocal microscopy, Brilliant Blue 515 (BB515)-conjugated goat anti–mouse immunoglobulin G (IgG) (Biolegend), Brilliant violet 421 (BV421)-conjugated goat anti-mouse IgG (Biolegend), Alexa633-conjugated donkey anti-goat IgG (ThermoFisher Scientific) for flow cytometry and confocal microscopy, streptavidin-HRP for WB (GE Healthcare, Piscataway, NJ) and anti-α_v_ integrin mouse mAb 272-17E6 (Sigma-Aldrich) for inhibition of TGF-β1 activation. Monoclonal antibody (clone 1/8.8) to human LRRC33 was described (*18*). For flow cytometry assay, anti-LRRC33 Ab was labeled by fluorophore Alexa-488 NHS Ester (ThermoFisher Scientific) as per the manufacturer’s instructions. Alexa488 conjugated antibody was purified with extensive dialysis in PBS buffer.

### Flow cytometry

PMA stimulated or non-stimulated MV4-11 and AML193 Cells were stained and analyzed as described previously (*10*). In brief, cells were resuspended and incubated with primary antibodies or direct conjugated antibodies at 1μg/ml in flow cytometry buffer (PBS with 2% FBS) for 60 min on ice. After washing with flow cytometry buffer cells were incubated with secondary antibodies for 30 min on ice and wash twice and analyzed by FACScanto instrument (BD Biosciences). Cell fixation and permeabilization was carried out using the Fixation/Permeabilization Solution Kit (BD Biosciences). The flow cytometry data was analyzed by Flowjo software.

### Confocal microscopy

AML193 and MV4-11 cells (with or without PMA stimulation) were harvested from cell culture and fixed with 4% paraformaldehyde (PFA) on glass slides treated with 0.1 mg/ml L-lysine. Fixed cells were treated with permeabilization and blocking solution containing 5% BSA, 2% FBS and 0.03% Triton X-100 (Sigma-Aldrich). Primary and secondary antibodies were diluted in PBS containing 5% BSA and 0.05% Triton X-100. Cells were incubated with the primary antibody (1μg/ml) overnight at 4 °C, washed with PBS, and incubated with the secondary Alexa fluorophore-conjugated antibodies (0.5μg/ml) for 1h at 4°C while protected from light. Slides were then washed with PBS, stained with DAPI for 5 min in room temperature, and coverslipped in ProLong Gold antifade reagent (Invitrogen) and imaged using Olympus Fluoview FV1000 confocal laser scanning microscope. LRRC33 was stained with anti-LRRC33 Ab and Alexa488-anti-mouse Ab; proTGF-β1 was stained with goat anti-LAP polyclonal Ab and Alexa633-anti-goat Ab. Non-specific mouse and goat IgG are used as primary Abs as controls.

### Western Blot (WB) and Immunoprecipitation (IP)

AML193 and MV4-11 cells (with or without PMA stimulation) were lysed with lysis buffer (25mM Tris, 150mM NaCl, 1mM EDTA, 1% NP-40, 5% Glyceral) with proteinase inhibitor cocktail (Roche) on ice for 30 min. The lysate was clarified by centrifugation at 12,000 × g for 5 min at 4°C and the lysate was applied to SDS-PAGE and analyzed by WB. IP is carried out with clarified cell lysate pre-cleaned by protein G sepharose (GE Healthcare), then the cell lysate was incubated with anti-LRRC33 or control mouse IgG (2μg/ml) overnight at 4°C on a rotator. Protein G Sepharose was then added and incubated at 4°C for another 1 h. The Sepharose beads were sedimented and washed three times with IP lysis buffer. The bound proteins were eluted by heating at 95°C in SDS sample buffer, separated by SDS–PAGE, and immunoblotted with antibody to prodomain. Data shown are representative of at least 3 independent assays.

### TGF-β1 bioassay

The TGFβ reporter cell line TMLC was a kind gift of Daniel Rifkin (New York University). The TGF-β1 activation assay was performed as previously described (*19*). In brief, AML193 and MV4-11 cells with PMA stimulated for 24h or without stimulation are washed twice with PBS and re-suspended in DMEM medium with 0.1%BSA before co-cultured with TMLC cells. Cells were co-cultured in 96-well plate overnight at 37 C° with 15,000 TMLC cells in each well and increasing numbers of MV4-11 or AML193 cells in DMEM + 0.1%BSA. For integrin Ab inhibition assay, 15,000 TMLC cells were co-cultured with 50,000 MV4-11 or AML193 cells in the presence of 2.5 μg/ml anti-α_v_ integrin (mouse mAb 272-17E6) or control mouse IgG. After co-culture, the cells were washed, lysed and processed using the Luciferase Assay System (Promega) and analyzed by Microplate Reader (BioTek). Data are presented as the mean and standard deviation of four replicates.

## Results

### Lrrc33 gene expression is the highest in AML cell lines

By searching on the publicly available database Cancer Cell Line Encyclopedia (CCLE), we identified that lrrc33 mRNA level is the highest in 39 AML cell lines compared to cell lines of other types of cancers (Fig. 1A). Furthermore, the mRNA level of LRRC33 and TGF-β1 are both high and correlated in the AML cell lines (Fig. 1B). Based on these data, we used two AML cell lines, MV4-11 and AML193, to investigate LRRC33 interaction with pro-TGFb1.

**Figure 1.**
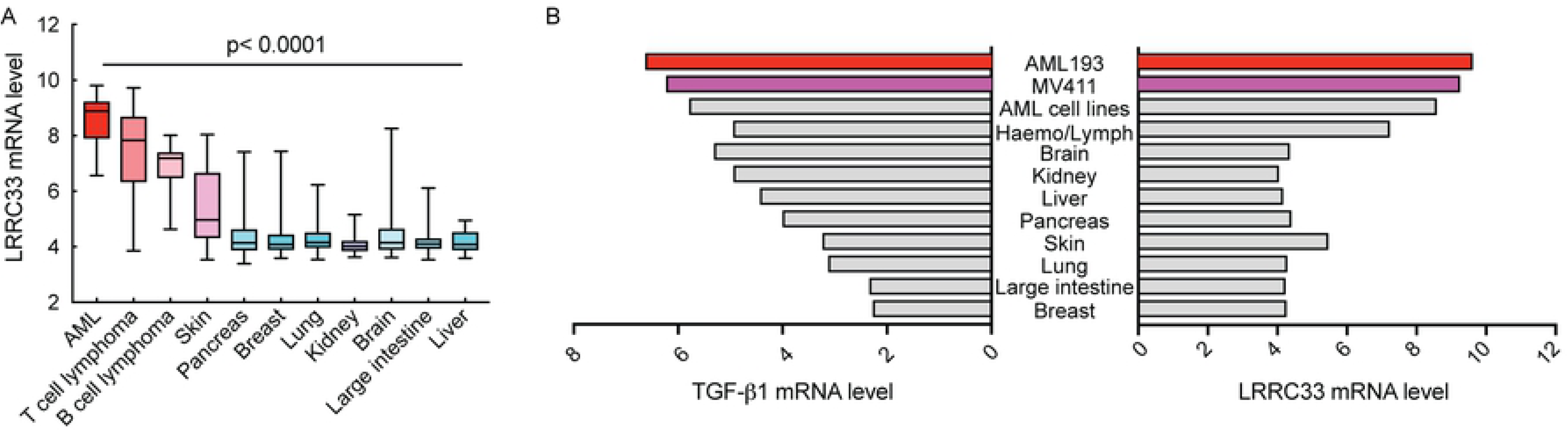
LRRC33 and TGFβ1 mRNA expression in acute myeloid leukemia (AML) and other cancer cells. Data is from Cancer Cell Line Encyclopedia database. (A) LRRC33 has the highest mRNA level in AML cells compared to other cancer cells. The p values are calculated by comparing mRNA level of AML cells to other cells through one-way analysis of variance (ANOVA) in Prism 7 Graphpad software. Error bars indicate the range of mRNA level of all the cell lines in each category. (B) LRRC33 and TGFβ1 mRNA expression are both high in AML cells. AML193 and MV4-11 are two representative cell lines studied in this research. Haemo/Lympo indicates other hematopoietic and lymphatic cells.

### LRRC33 and pro-TGF-β1 are expressed on the surface of MV4-11 and AML193 cells, and the protein level increases after PMA stimulation

To investigate the location of LRRC33 and pro-TGF-β1 in MV4-11 and AML193 cells, we carried out flow cytometry experiments using LRRC33 and pro-TGF-β1 mAbs. The Cell surface staining showed that both LRRC33 and pro-TGF-β1 were expressed on the cell surface of MV4-11 and AML193 cells, and the expression level of both proteins were increased after PMA stimulation for 24 hours (Fig. 2A). PMA is utilized as stimulus of cytokine synthesis in the cancer cell lines. Staining of fixed and permeabilized cells indicated that the total amount of protein expression of LRRC33 and pro-TGF-β1 also increased significantly after PMA stimulation (Fig. 2B). These results indicated the expression of LRRC33 and pro-TGF-β1 was correlated in both non-stimulated and stimulated cells, and suggested that LRRC33, similar to GARP in Treg cells (*9*), is a potential pro-TGF-β1 anchoring protein in the MV4-11 and AML193 cells for TGF-β1 activation.

**Figure 2.**
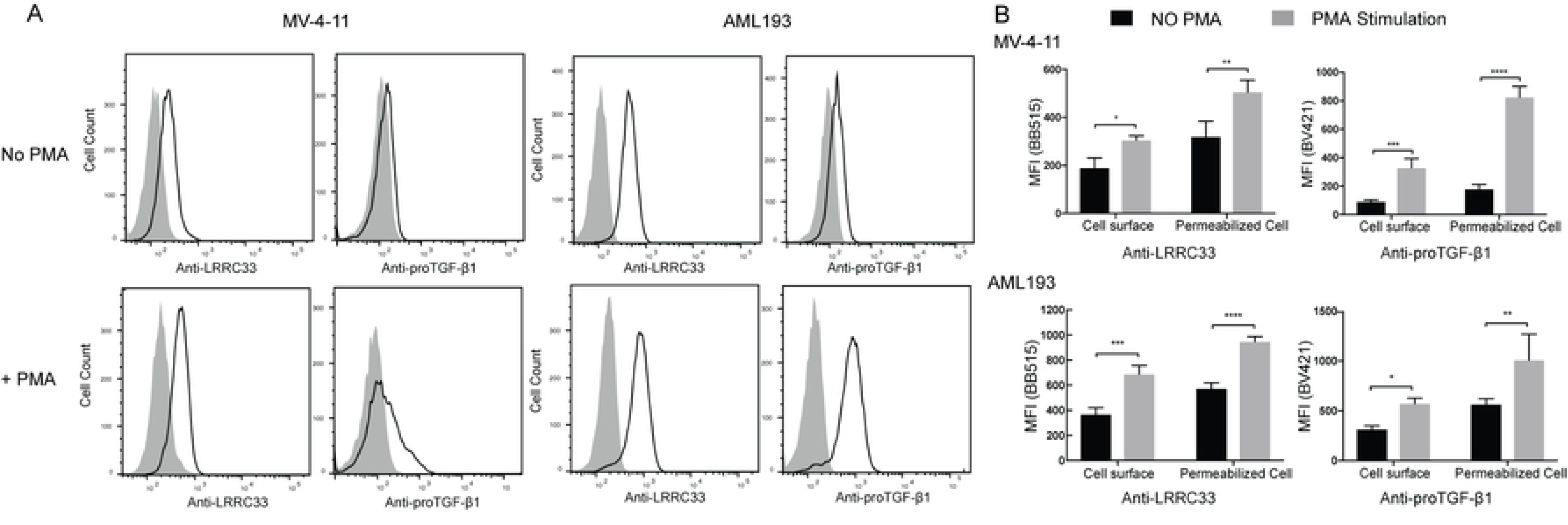
Expression of LRRC33 and pro-TGF-β1 proteins in MV4-11 and AML193 cells by flow cytometry. Cells were stained with anti-LRRC33 and anti-pro-TGFβ1 antibodies (Abs), and mouse IgG isotype control was used as primary antibody in control staining. BB515 or BV421-conjugated goat anti–mouse immunoglobulin G (IgG) was used as secondary Abs. Data are representative of three independent experiments. (A) Cell surface staining of LRRC33 and pro-TGF-β1 with or without PMA stimulation. The shaded curves indicate the IgG control. (B) Comparison of total LRRC33 and pro-TGF-β1 protein expression with or without PMA stimulation. Cells were permeabilized before staining. The amount of proteins is indicated by the mean fluorescence intensity (MFI). The background MFI from control IgG staining was subtracted from each group. Error bars represent the standard deviation of 3 replicates. The p values are calculated by 2-way ANOVA comparisons in Prism 7 Graphpad software. ns: not significant. *: p<0.1, **: p<0.01, ***: p<0.001, ****: p<0.0001

### LRRC33 and pro-TGF-β1 form complex and co-localize on the cell surface in the MV4-11 and AML193 cells

To investigate whether LRRC33 and pro-TGF-β1 are co-expressed in MV4-11 and AML193 cells, flow cytometry with double staining using anti-LRRC33 and anti-pro-TGF-β1 mAbs with fixed and permeabilized cells was performed. LRRC33 was expressed on almost all the cells with or without PMA stimulation, while there are 38.5% of MV4-11 and 74.1% AML193 cells expressing both proteins without PMA stimulation and 84.5% of MV4-11 and 95.1% AML193 cells expressing both proteins after PMA stimulation (Fig. 3A). These data indicated that there is no sub-group of cells that only express pro-TGF-β1 without LRRC33 and most of the stimulated cells expressed LRRC33 and pro-TGF-β1 at the same time. Further, co-localization of LRRC33 and pro-TGF-β1 was observed through confocal microscopy. Fig 3B shows that after PMA stimulation, LRRC33 co-localized with pro-TGF-β1 on the cell surface forming a ring-shaped pattern and in the plasma outside the nucleus (Fig. 3B). The co-localization is not significant in non-PMA stimulated cells, due to low pro-TGF-β1 protein expression without PMA stimulation (Fig. 3A, 3B). We further investigate if LRRC33 interacts with pro-TGF-β1 to form protein complex using immunoprecipitation (IP) and Western blotting (WB). With or without PMA stimulation, MV4-11 and AML193 cell lysates were blotted with antibody to the TGF-β1 prodomain, or IP with the anti-LRRC33 mAb followed by blotting with anti-pro-TGF-β1 Ab. The WB of cell lysate showed that without PMA stimulation, the expression level of pro-TGF-β1 and protein complex is low in MV4-11 cells, whereas pro-TGF-β1 and protein complex level are higher in AML193 cell, possibly due to different background cytokine activation level in those two cell lines (Fig. 4A). After PMA stimulation expression of both pro-TGF-β1 and the complex increased in both cells with a more significant increase of pro-TGF-β1 alone (Fig. 4A). We further indicate the protein complex observed in WB is the LRRC33-pro-TGF-β1complex by IP experiment. The IP with anti-LRRC33 mAb and blotting with anti-pro-TGF-β1 Ab demonstrated the association of LRRC33 with pro-TGF-β1 and an increase of protein complex formation after PMA stimulation (Fig. 4B). The detection of pro-TGF-β1 band in reducing SDS-PAGE compared to the protein complex band in the non-reducing SDS-PAGE indicated that LRRC33 interacts with pro-TGF-β1 through disulfide links (Fig. 4B). These results demonstrated that LRCC33 binds and disulfide links to pro-TGF-β1 and provided further evidence of LRRC33 being a pro-TGF-β1 regulating protein.

**Figure 3.**
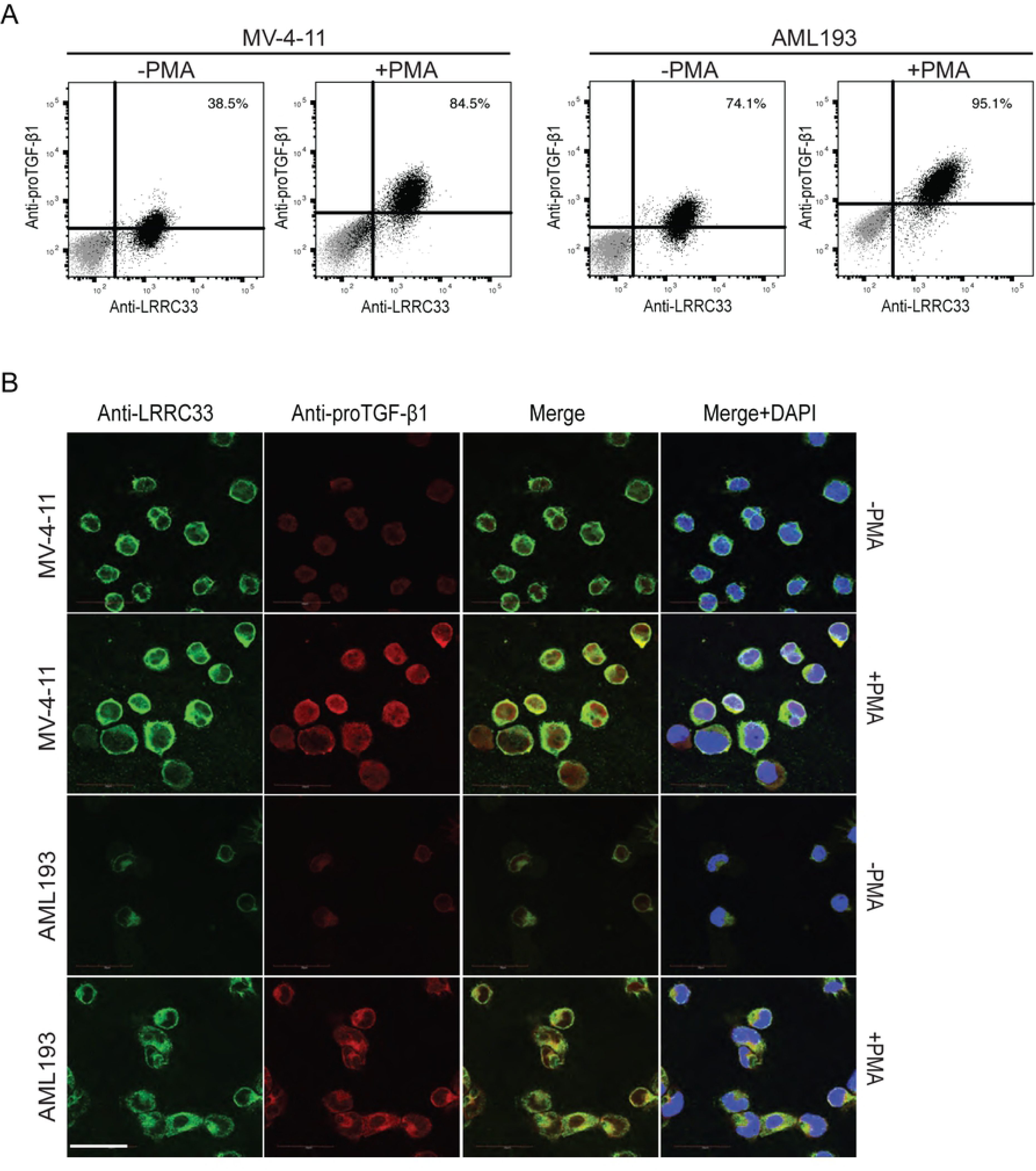
LRRC33 co-localizes with pro-TGF-β1 in MV4-11 and AML193 cells. (A) Flow cytometry of double-staining of fixed and permeabilized AML193 and MV4-11 cells. Cells were stained with Alexa488-conjugated LRRC33 and APC-conjugated pro-TGF-β1 mAbs before and after PMA stimulation. Alexa488- and APC-directly conjugated non-specific IgG were used as control and the control staining are indicated in grey dots. Numbers of double positive cells are shown. (B) LRRC33 and pro-TGF-β1 co-localization visualized by confocal microscopy. Green: anti-LRRC33; Red: anti-pro-TGF-β1; Blue: DAPI staining.

**Figure 4.**
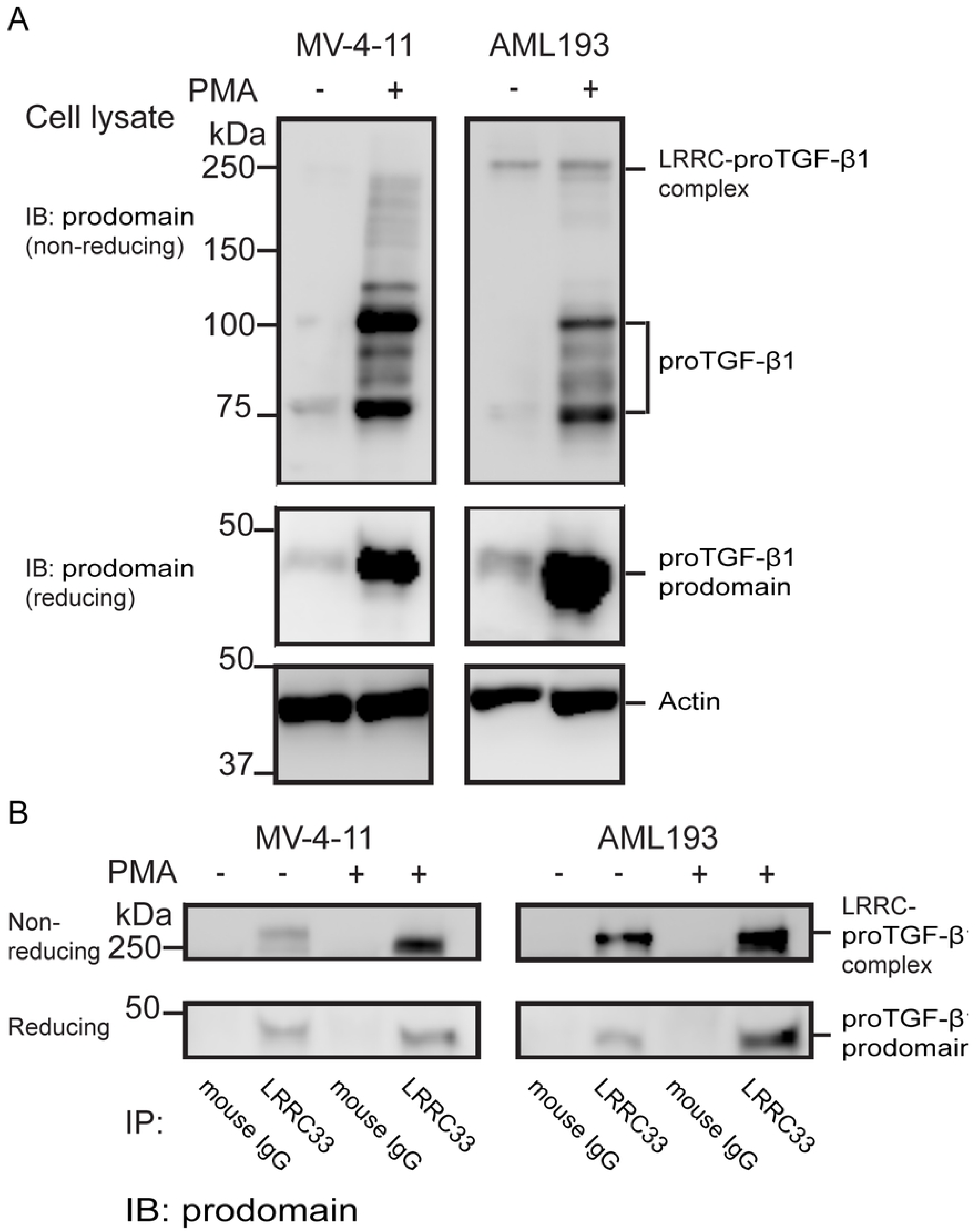
Detection of LRRC33-pro-TGF-b1 complex in MV4-11 and AML193 cells by Western blot (WB) and immunoprecipitation (IP). (A) Cell lysates were analyzed by immune-blotting using biotinylated pro-TGF-β1 Ab. The LRRC33-pro-TGF-β1 complex band is indicated in non-reducing SDS-PAGE, and the band of pro-TGF-β1 alone is indicated in both non-reducing and reducing SDS-PAGE. (B) Co-immunoprecipitation. Cell lysates were subjected to immunoprecipitation using LRRC33 antibody or control mouse IgG as indicated, and the immunocomplexes were analyzed by WB using pro-domain antibody. The immunoprecipitated LRRC33-pro-TGF-β1 complex is indicated in non-reducing SDS-PAGE, and the pro-TGF-β1 pull down is indicated in the reducing SDS-PAGE.

### TGF-β1 activation in MV4-11 and AML193 cells was integrin dependent

We next studied TGF-β1 activation in MV4-11 and AML193 cells with or without PMA stimulation using mink lung TGFβ-reporter cell line (TMLC) system. The α_V_β6 and α_V_β8 integrins have been shown to activate TGF-β1 (*20, 21*). Our result showed that the level of TGF-β1 activation increased in MV4-11 and AML193 cells after PMA stimulation (Fig. 5A and 5B), and such activation was inhibited by adding anti-α_V_ integrin Ab to the PMA stimulated AML193 and MV4-11 cells cultured with TMLC cells (Fig. 5C). This result indicated that TGF-β1 activation in MV4-11 and AML193 cells was α_V_ integrin-dependent, similar to other cells expressing LTBP or GARP as pro-TGF-β1 ligand. These results further confirmed our hypothesis that LRRC33 is a cell surface binding partner of pro-TGF-β1 and regulate the activation of TGF-β1 through α_V_ integrin-dependent mechanism like other pro-TGF-β1 binding proteins.

**Figure 5.**
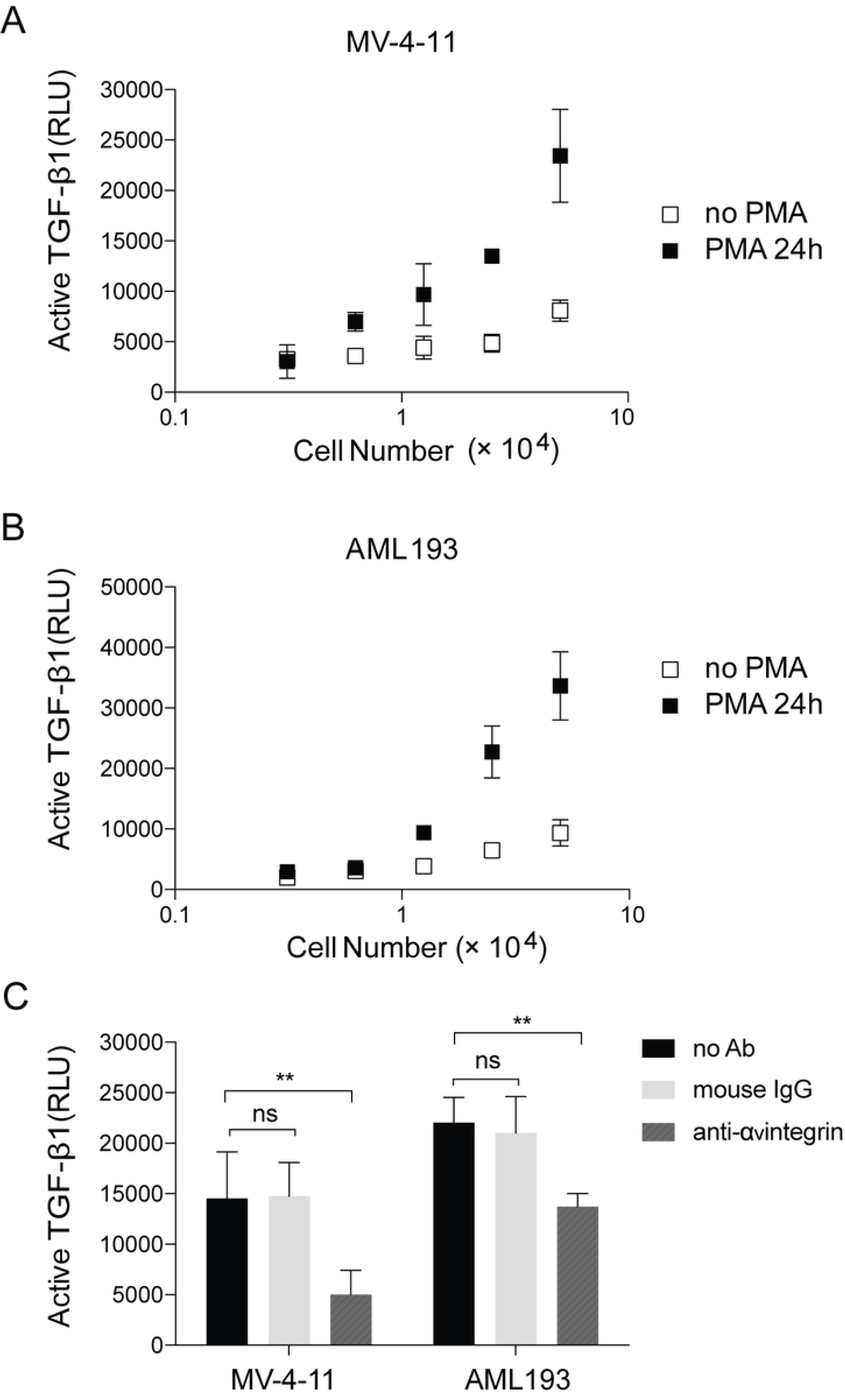
TGF-β1 activation in MV4-11 and AML193 cells assessed in TMLC-luciferase reporting system. The luciferase activity is expressed in relative light units (RLU). Data are presented as the mean and standard deviation of four replicates and is the representative of three independent experiments on different days. (A) MV4-11and (B) AML193 cells, without or with PMA stimulation. (C) For MV4-11 and AML193 cells with PMA stimulation, incubation with 2.5 μg/ml anti-α_v_ integrin mouse mAb 272-17E6 results in decrease of TGF-β1 activation. Non-specific mouse IgG is used as negative control. The p values are calculated by 2-way ANOVA comparisons in Prism 7 Graphpad software. ns: not significant. *: p<0.1, **: p<0.01, ***: p<0.001, ****: p<0.0001

## Discussion

In this study, we demonstrated that a recently identified pro-TGF-β1 binding protein, LRRC33, associated with pro-TGF-β1 in human myeloid leukemia cell lines and played a role in TGFb1 activation in these cells. In two models of human myeloid leukemia cell lines, MV4-11 and AML193, LRRC33 co-localizes and forms protein complex with pro-TGF-β1. Although protein expression of pro-TGF-β1 largely relies on activation by PMA, the LRRC33 has high level of background expression without PMA stimulation (Fig. 2, 3 and 4), indicating a potential housekeeping role of LRRC33 in AML cells. We also show that TGF-β1 activation in these two cells lines is integrin dependent like LTBP and GARP expressing cells, indicating LRRC33 is the regulating protein of TGF-β1 function in human myeloid leukemia cells.

In previous researches, LRRC33 is known as the Negative Regulator of Reactive Oxygen Species (NRROS), and its function is down-regulating of reactive oxygen species during host defense and autoimmunity on gene knock-out mice (*22*). And a following research by another group indicates that LRRC33 regulates osteoclast differentiation (*23*). In our previous research, it is indicated that the LRRC33 homologous protein, GARP (LRRC32), binds to pro-TGF-β1 and exert TGF-β1 regulating functions (*17*). Further, our recent publication demonstrates that LRRC33 is expressed in microglia cells in the CNS and Lrrc33 knockout mice appear to develop CNS vascular abnormalities associated with deficiency in TGF-β activation (*18*). These evidences prompt our hypothesis of LRRC33 as a novel binding and regulating protein of TGF-β1 in myeloid lineage cells. Nevertheless, our findings are not necessarily contradictory to the aforementioned studies given that TGF-β1 has been shown to negatively regulate innate immunity pathways (*24*) and down regulate both secretion of pro-inflammatory cytokines and reactive oxygen species production (*25*).

The mRNA array data of LRRC33 and TGF-β1 from CCLE (Fig. 1) indicate LRRC33 could play pivotal roles in AMLs in the context of TGF-β signaling pathway. It has been reported that treatment of THP-1, another human myeloid cell derived leukemia, with TGF-β1 reduces the growth rate of cells and promotes the differentiation to macrophage (*26*). Another report show that TGF-β1 is able to induce the differentiation of human myelogenous leukemia cell line U-937 into macrophages (*27*). In addition, other chemical differentiation factors including PMA and retinoic acid were also indicated to induce cell differentiation of promyelocytic leukemia cells HL-60 into macrophage (*15*). TGF-β1, PMA and retinoic acid were suggested to be potential treatment for myeloid cell derived leukemia in these researches. However, in our research we indicate that PMA stimulation of MV4-11 and AML193 cells result in significant increase of the expression level of TGF-β1 and LRRC33. In the microenvironment of multiple malignancies TGF-β1 has immune suppressive function, inhibiting the activation and proliferation of T cell, B cells and natural killer cells (*28-30*). In addition, acute myeloid leukemia cells exhibit resistance to TGF-β1 growth inhibition effect in the process of tumor progression (*14*). Thus, the stimulation of myeloid leukemia cell may cause suppression of immune cell function due to excessive TGF-β1 production. Further, in clinical practice, the complex genomic landscape of AML makes it difficult to develop widely applicable targeted therapy. Taken together, LRRC33 or LRRC33-pro-TGF-β1 complex is a potential myeloid cell specific target to regulate TGF-β1 function in cancer cells and tumor microenvironment for non-genomic based therapy of AML and other myeloid cell malignancy in a broader context.

